# Geographical origin and cultivar differentiation of Kava (*Piper methysticum*) using Artificial Neural Network with FTIR Spectroscopy: A Novel Method

**DOI:** 10.1101/2025.10.05.679113

**Authors:** Ronick Spenly Shadrack, Daniel Tari, Hancy Tabi, Jacinta Botleng, Rolina Kelep, Ladyshia Regenvanu, Cassandra Tangaras, Mowa Pakoasongi, Edword Butjukabwaelep, Galana Siro, Atanas Pipite, Vincent Lebot

## Abstract

This study presents a novel method for authenticating the geographical origin and cultivar of kava (*Piper methysticum*) by combining Fourier Transform Infrared (FTIR) spectroscopy with Artificial Neural Networks (ANN). A spectral database of kava varieties from four (4) countries in the Pacific Island region, namely Vanuatu, Fiji, Papua New Guinea, and Hawaii, was used for regional authentication. For samples collected within Vanuatu, spectral data were obtained from the acetone extract of both fresh and dried kava. The ANN predictive model was trained on geographical origin (countries or islands of origin), quality (noble vs tudei), and between different cultivars. ANN achieved near-perfect performance, with a generalized R-Square of 0.99 (training), 0.84 (validation), and 0.95 (test) for geographical origin prediction. Class-specific accuracy was 100% for Vanuatu, Papua New Guinea, and Hawaii. The model achieved near-perfect internal prediction accuracy (R² = 0.99) under repeated-measure validation. Due to the limited availability of independent Fiji samples, no blind-test validation was performed for that region. Thus, robustness claims apply only to regions with sufficient validation data.

Significantly, the model demonstrated perfect classification (100% accuracy) for Malo and Santo Island kava samples, highlighting its ability to authenticate micro-regional origins within Vanuatu. For variety differentiation, ANN achieved 100% accuracy for noble versus tudei cultivars, ensuring compliance with Vanuatu’s noble-only export policy. ATR-FTIR spectra of fresh and dried kava acetone extracts exhibited visually distinct patterns among kava cultivars at spectral regions of 1750 cm^-1^ to 1525 cm^-1^ and 1124 cm^-1^ to 900 cm^-1^, indicating potential for direct differentiation. Visual detection of kava adulteration at 1585 cm^-1^ was feasible at 1% tudei or wichmannii substitution without advanced analysis. These findings position ANN-FTIR as a rapid, non-destructive, and cost-effective solution for food authentication, geographical indication labeling, and export certification, supporting international standards such as Codex Alimentarius and International Standards Organization (ISO) guidelines.

**Graphical abstract:** 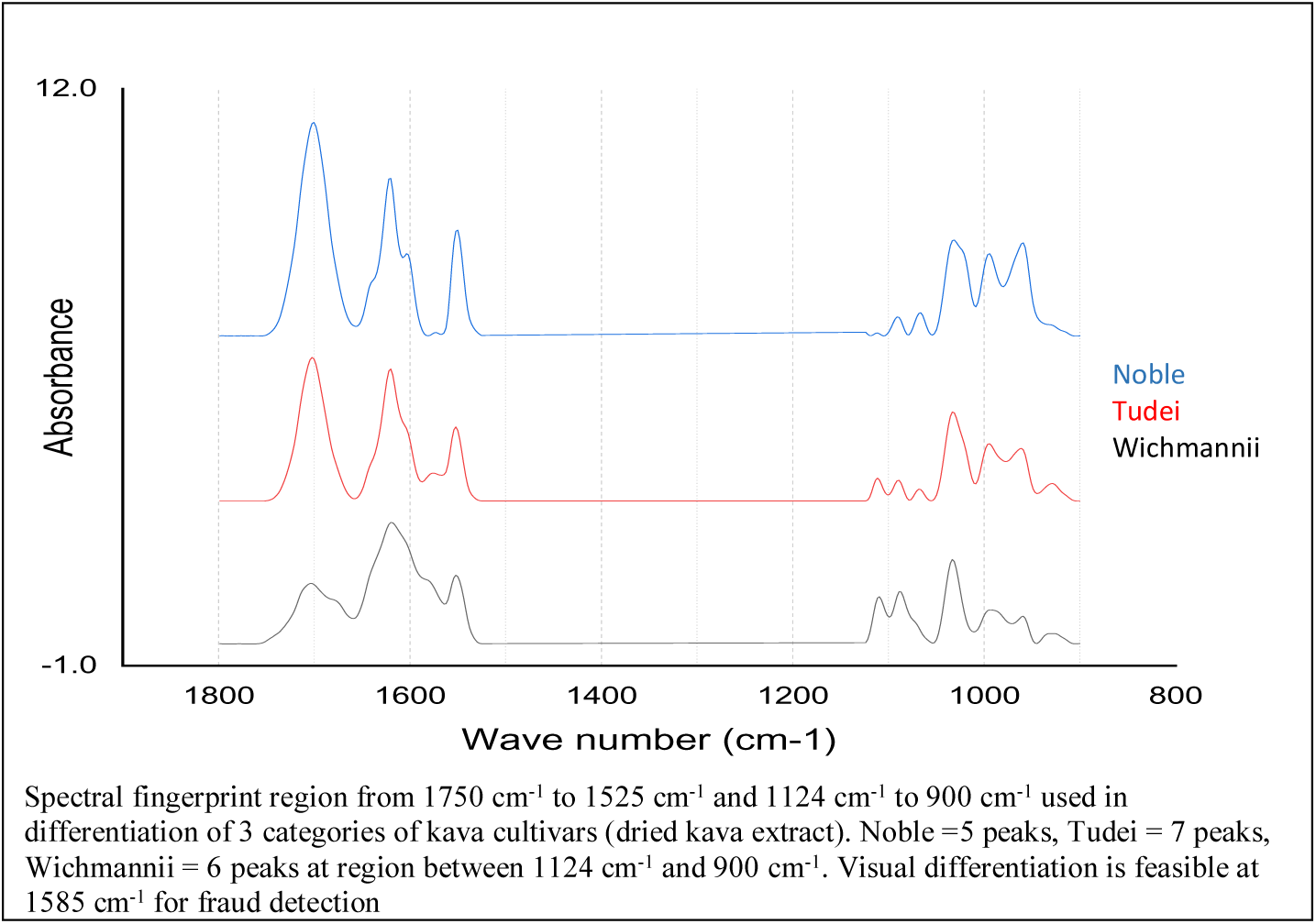

## 1. Introduction

Kava (*Piper methysticum*), a key economic crop of the Pacific Islands, is widely consumed as a beverage and increasingly marketed as a dietary supplement. Its commercial value depends on both geographical origin and varietal classification, with noble kava being preferred for export due to its desirable pharmacological properties and lower toxicity compared to tudei kava (1, 2).

Despite its economic importance, kava authentication remains challenging. Conventional approaches, including genetic analyses (DNA barcoding) and chromatographic profiling (HPTLC, HPLC or LC-MS), are effective but are time-consuming, costly, and require specialized laboratory facilities. These constraints limit routine screening of large sample volumes needed for export certification and regulatory compliance. Moreover, differentiating kava at the micro-regional level, such as between islands within Vanuatu (e.g., Malo vs. Santo), has been difficult due to overlapping chemical profiles and environmental variability (3, 4). Different statistical models were used with chemometric data to predict kava quality with near-infrared (IR) spectroscopy (5). Traditional linear models such as principal component analysis (PCA), linear discriminant analysis (LDA), partial least squares discriminant analysis (PLS-DA), and soft independent modelling of class analogy (SIMCA) have proven useful for classification and exploratory analysis. However, their performance is limited by the inherent complexity and nonlinear nature of the kava chemical matrix. To address this, nonlinear modeling approaches such as artificial neural networks (ANN), support vector machines (SVM), and random forests (RF) may provide improved predictive accuracy in the assessment of kava quality (6-8). These methods are capable of modeling complex, nonlinear interactions among chemical constituents, which are often not captured by linear methods. While non-linear statistical approaches enhance prediction accuracy in kava quality assessment, the effectiveness of this approach is heavily dependent on input data which captures the chemical properties or complexities of kava. Fourier Transform Infrared (FTIR) Spectroscopy, provides comprehensive spectral fingerprints making it a profound technique when coupled with non-linear models. Hence, combining FTIR spectroscopy with chemometric or machine learning models emerged as a promising solution for rapid, non-destructive authentication of food and herbal products, consequently, optimizing daily operations or routine work. Several empirical studies have been recently developed incorporating non-linear approaches with FTIR such as analysis of soil organic carbon (9), classification of honey (10), milk authentication (11), differentiating Gram-negative and Gram-positive bacteria (12), and evaluation of contaminants in wheat flour (13). Specifically, FTIR captures the overall chemical fingerprint of plant material, allowing subtle differences in chemical composition to be detected. When coupled with Artificial Neural Networks (ANNs), which can model complex, non-linear relationships in spectral data, this approach offers highly accurate classification even in challenging cases (14, 15).

While recent researches have demonstrated the veracity of coupling machine learning models with FTIR in authenticating diverse commodities, its application to kava quality assessment remains largely unexplored. Implementing such methods for kava authentication carries several important implications. First, it enhances quality control by ensuring that exported kava adheres to noble-only policies, thereby protecting consumers from the potential toxicity of tudei cultivars. Second, it strengthens traceability and fraud prevention by enabling verification of geographical origin down to individual islands, which supports geographical indication labeling and adds value through premium marketing. Third, it reduces the time required for analysis and facilitate fast trading of goods between regions. Finally, it boosts market confidence by reinforcing international trust and facilitating compliance with regulatory standards, including Codex Alimentarius and ISO 22000, which are essential for participation in the global trade network (16).

Given these motivations, this study presents a novel ANN–FTIR methodology designed for comprehensive kava authentication. It evaluated kava based on: (i) geographical origin across multiple counties in the Pacific Island region, (ii) differentiation between noble and tudei cultivars, (iii) classification based on dried and fresh extracts, including micro-regional island-level differentiation (Malo and Santo island of origin), and (iv) differentiate and detect fraud in samples of noble kava with tudei cultivars. This integrated approach offers a rapid, accurate, and scalable tool for kava quality control, regulatory compliance, and export certification with aims to enhance authenticity verification processes, support regulatory frameworks, and promote consumer confidence in kava products.

## 2. Methods

### 2.1 Sample Collection

A total of 150 kava samples, comprising noble and two day (tudei) cultivars, were collected from multiple locations across Malo and Santo Island in Vanuatu. The samples were authenticated by experienced farmers and verified based on morphological and chemo-type characteristics (18). Based on the knowledge of kava cultivars, most recent local common names were used such as green hand (melomelo), red hand (borogu), yellow leave (palarasul), ambae (bilitudei), palisi (tudei), klish hand (tudei), wild kava or flower kava (wichmannii) with other less common cultivars in the two islands. All these names and quality were verified with chemical finger printing and multivariate analysis. For samples category of kava plants, samples were categorized into 3 subgroups, the roots, chips and unpeeled chips and data were collected according to each of these fraction.

### 2.2 Sample Preparation

Fresh kava stem and root samples were peeled or washed to remove residual soils and debris, chopped, and dried in a dehydrator at 60 °C until constant weight was achieved. The dried material was grounded to a uniform particle size using a dried food (KA3025, China) grinder and subsequently sieved. Less than 0.5 g of the powdered sample was placed directly onto the attenuated total reflectance (ATR) diamond crystal of Brukers Alpha II FTIR spectrophotometry (Germany) for spectral collection. Prior to spectral collection, fresh kava samples pulverised and stored at room temperature for acetone extraction.

### 2.3 Kava Lactone Extracts

Extracts were obtained from freshly minced and oven-dried powdered kava samples using acetonitrile in a 3:1 w/w ratio, following the method of Lebot et al. (2). The extracts were then evaporated in a dehydrator at 50 °C for 12 hours. The resulting lactone paste transferred using a stainless-steel micro spatula (149 mm length, 5 mm flat spade tip) onto the ATR diamond of Brukers Alpha II FTIR crystal for spectral analysis. For the detection of tudei and wichmannii kava adulteration in noble kava samples, pulverized kava samples, pre-analysed for baseline composition, were used for detection of tudei cultivar following the method of Lhuissier et al. (19) with modifications. For adulteration, mixtures of noble kava containing varying percentage proportions of tudei and wichmannii kava were prepared at the following ration: 0, 0.5, 1, 2, 3.5, 5, 10, 20, 30, 40, 50, 60, 70, and 100 percent proportion. Each mixture was extracted from a 1:3 (kava: acetone) solution and the extracts were centrifuged at 10,000rpm. The supernatant was collected and measured using a colorimeter at wavelength of 440 nm. The solution extracts were transferred to 10 ml glass test tubes, and acetone was removed by dehydrating the solution extract at 50°C for 12h. The dried extracts were stored at room conditions for continual analysis with FTIR spectrophotometry.

### 2.4 FTIR Spectroscopy

FTIR spectra were recorded in the range of 4000–400 cm⁻¹, with 2 cm⁻¹ resolution and 23 scans averaged per sample. The regions from 1800 cm^-1^ to 900 cm^-1^ was used in the analysis. All spectra were baseline-corrected, normalized, smoothed and subjected to standard normal variant corrected (SNV) prior to analysis. For regional kava samples, the spectral database of 138 kava samples was obtained from Lebot et al., (5). Originally, this dataset was used with PCA, LDA and PLS-DA analysis. In this study, the same database was re-analysed with ANN for comparison. Briefly the spectra database was collected using Perkin Erlenmeyer FTIR (USA) spectrophotometer in the range of 1800 – 800 cm^-1^, with resolution of 1 cm^-1^ and 32 scans average per sample. The spectral dataset was baseline corrected, smooth and processed with first derivative.

### 2.5 Data Processing and ANN Development

The spectral dataset was randomly divided into training (60%), validation (25%), and blind test (15%) sets. A single hidden-layer ANN with three TanH neurons was trained using holdback validation (0.3), an additive sequence boosting algorithm (learning rate = 0.1), and a squared penalty fitting method over one boosting iteration.

For sample groups with small sample sizes, a single hidden-layer ANN with five TanH neurons was trained using 5-fold cross-validation, an additive sequence boosting algorithm (learning rate = 0.1), and a squared penalty fitting method over a single boosting iteration.

An artificial neural network (ANN) was implemented with JMP (USA) software, and its performance metrics were evaluated. The modelling approach followed previously established ANN applications in food authentication (20-22).

## 3. Results

### 3.1 FTIR Spectral Profile

The figure 1 presents the averaged FTIR spectra of dried kava powder samples derived from genetically characterized noble and tudei cultivars based on 138 between the two cultivars across the 4 island countries (Vanuatu, Fiji, Hawaii and PNG). The absorbance values span the mid-infrared fingerprint region (1800–800 cm⁻¹), which is rich in vibrational modes of organic functional groups relevant to plant secondary metabolite. The spectral differences align with known chemotypic distinctions between noble and tudei kava. Tudei samples typically contain higher levels of flavokavains (A, B, C) and related phenolic compounds, which exhibit characteristic absorbance bands in the aromatic and carbonyl regions. Noble cultivars, by contrast, show a more subdued spectral profile, consistent with lower concentrations of these compounds and a more balanced kava lactone profile.

**Figure 1.**
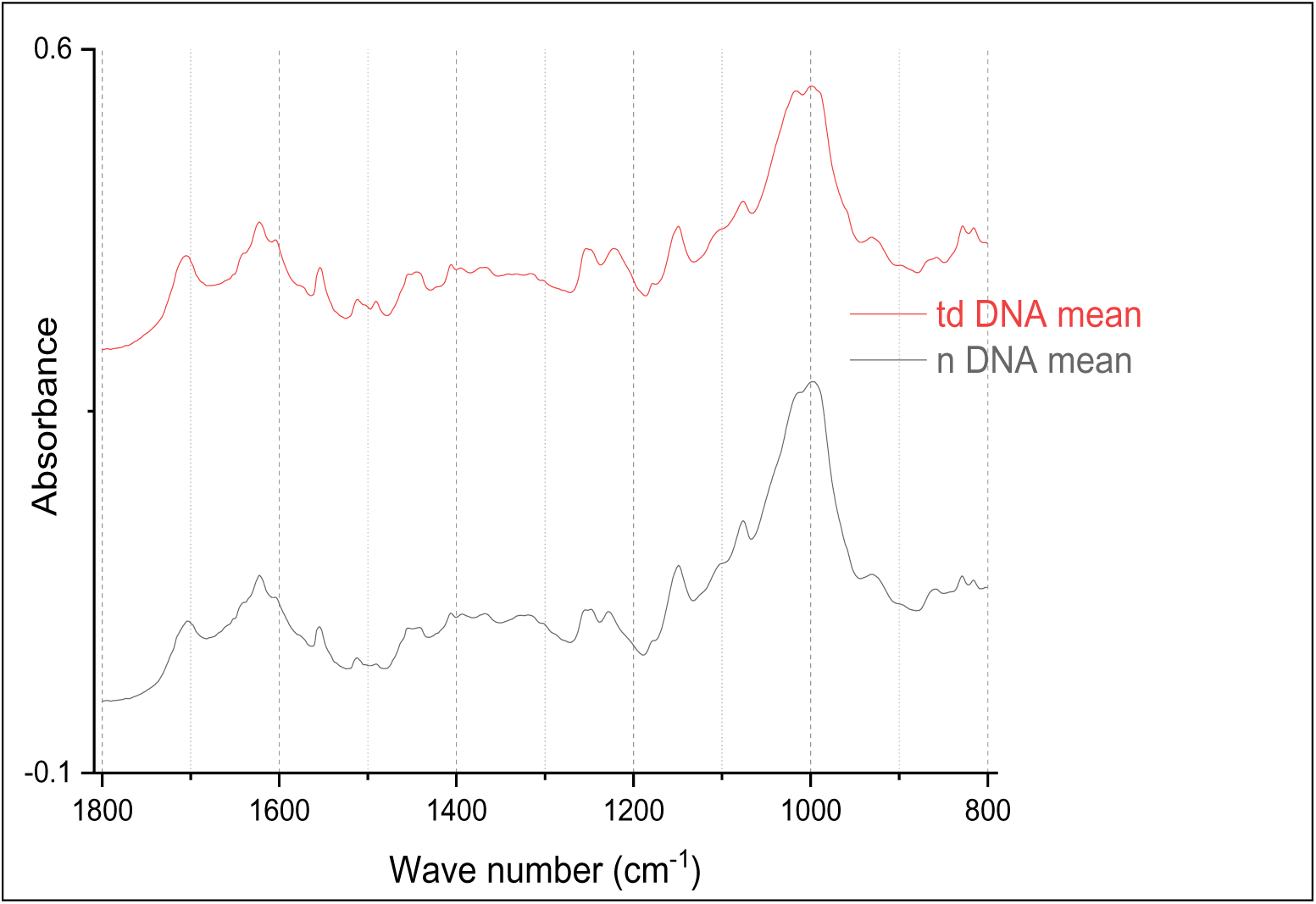
Average FTIR spectral data for dried kava samples in 4 countries classified with DNA finger printing (138 samples). The letter n represents noble and td represent tudei.

The classification of kava samples using fresh acetone extracts revealed clear spectral differences among cultivars (Figure 2). Noble cultivars (n = 90) exhibited spectra without a peak around 1100 cm⁻¹, whereas tudei and wichmannii cultivars (n = 38) displayed a prominent peak in this region, corresponding to dihydrokavain (DHK) and dihydromethysticin (DHM)

**Figure 2.**
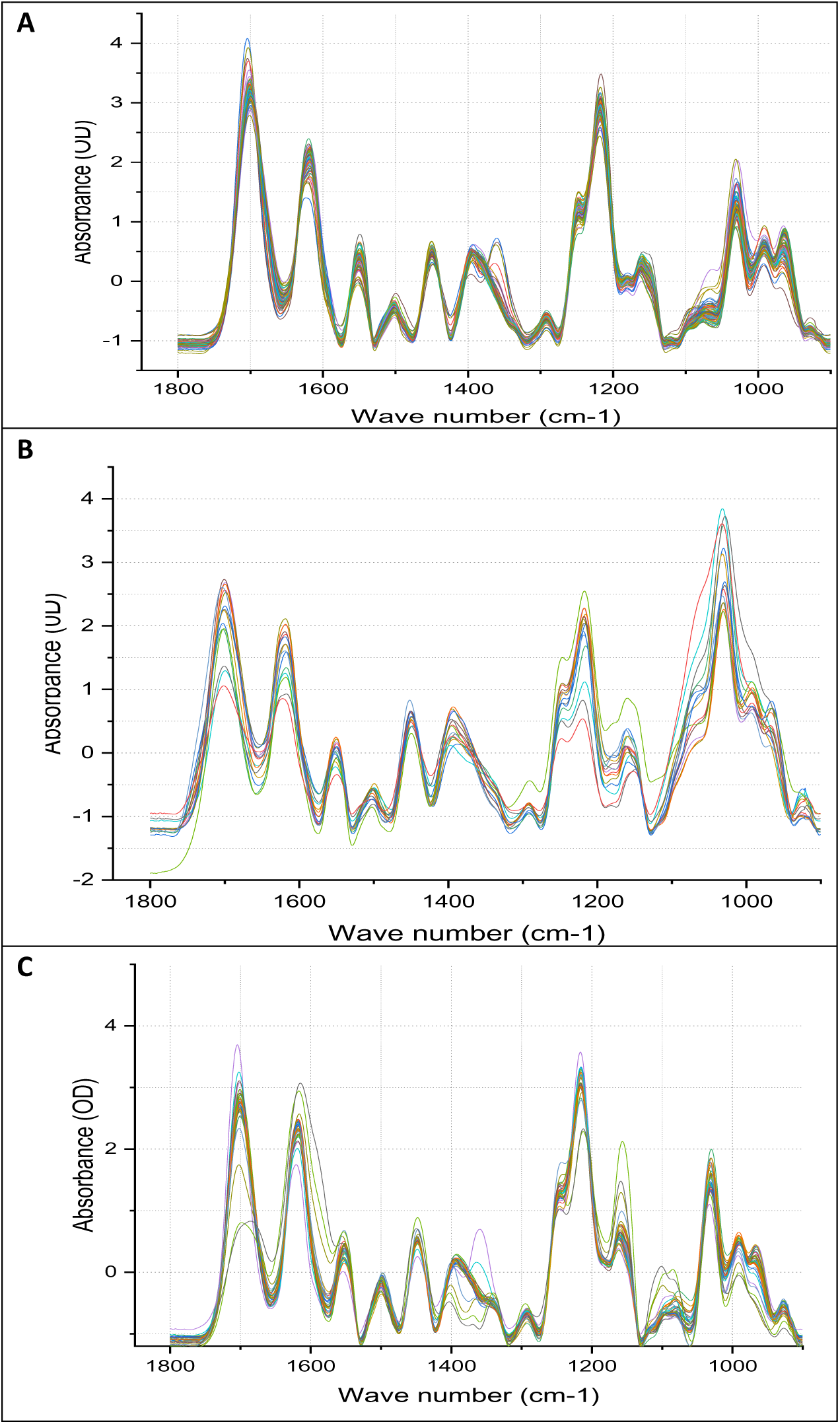
Spectral distribution of acetone extract of fresh kava. Samples collected from Malo and Santo, Vanuatu. A., Spectra of noble kava, B., Spectra of suspected tudei kava, C., Spectra of tudei and wichmannii kava.

(Table 1). Samples suspected of being tudei or mixed (n = 15) showed broad, high-intensity peaks spanning 1000–1100 cm⁻¹, indicating a potentially high concentration of DHK. These spectral distinctions demonstrate the utility of ATR-FTIR in differentiating kava cultivars based on their kavalactones composition.

**Table 1.** Kavalactones FTIR peak assignment and associated Kava cultivars.

| **Wavenumber (cm<sup>-1</sup>)** | **Functional Group** | **Dominant Kavalactone** | **Associated Kava Type** |
| --- | --- | --- | --- |
| **~1650** | C=O Stretch (Conjugated) | Dihydromethysticin (DHM) | Tudei / Wild |
| **~1650** | C=O Stretch (Conjugated) | Dihydrokavain (DHK) | Tudei / Wild |
| **~1600-1500** | C=C Aromatic | Yangonin / Desmethoxyyangonin | Noble |
| **~1450-1400** | C-H Bending | Kavain | Noble |
| **~1250-1000** | C-O Stretch / C-O-C | DHM DHK | Tudei / Wild |
| **~980-950** | C=C-H Out of plane | Methysticin | Noble |

The ATR-FTIR spectra of fresh acetone extracts revealed visually distinct patterns across kava categories, including noble, suspected tudei, tudei, and wichmannii (Figure. 3A). Noble kava was characterized by the absence of a peak at 1100 cm⁻¹. In contrast, suspected tudei exhibited a broad, high-intensity peak centred on 1050 cm⁻¹. Both tudei and wichmannii kava displayed peaks at 1100 cm⁻¹, with wichmannii further distinguished by a pronounced peak at 1625 cm⁻¹ and comparatively lower absorbance in the 1700 cm⁻¹ and 1550 cm⁻¹ regions when compared to other cultivars.

**Figure 3.**
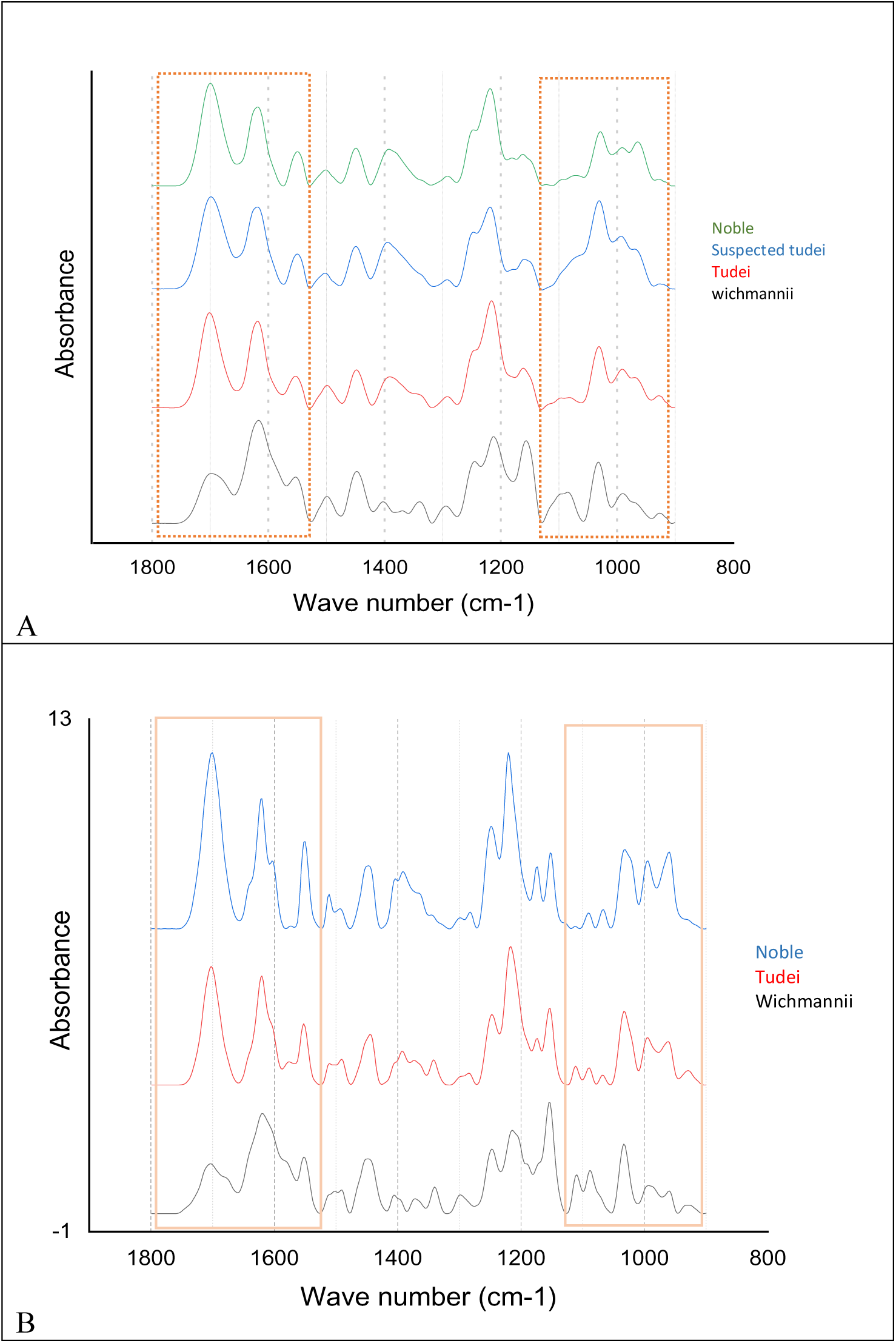
Distinct spectral differences between fresh and dried kava acetone extracts across noble, suspected tudei, tudei, and wichmannii cultivars. A., Spectra of fresh kava acetone extract, B., Spectra of dried kava acetone extract.

Figure 3B shows the FTIR spectra of dried kava acetone extracts, revealing distinct patterns across the three categories: noble, tudei, and wichmannii (wild kava). In the 1124 - 900 cm⁻¹ region, which corresponds mainly to C–O and C–N stretching vibrations, noble kava displayed five (5) distinct peaks, tudei exhibited seven (7), and wichmannii showed six (6). These variations in peak number and intensity highlight differences in the underlying chemical composition and provide a reliable spectral feature for discriminating between the categories. As shown in Fig. 3B, noble kava exhibit fewer peaks than tudei and wichmannii cultivars, this is attributed to the less substituted aromatic ring structure of kavain. Tudei and wichmannii kava exhibit more peaks, corresponding to the heavily substituted aromatic ring of dihydromethysticin and chemical composition of dihydrokavan. At 1625 cm^-1^, wichmannii showed higher peak intensity compared to other vareities which can be easily differentiated. The distinct features that differentiate all vareities lies in the region 1750 cm^-1^ to 1525 cm^-1^ and 1124 cm^-1^ to 900 cm^-1^. These region alone can differentiate cultivars in fresh and dried samples without need for machine learning algorithm.

The spectral distribution of dried kava sample acetone extract is shown in figure 4. The spectral peaks exhibit consistency with minimal variation, likely attributed to the standardised moisture removal procedure applied to the samples prior to spectra collection.

**Figure 4.**
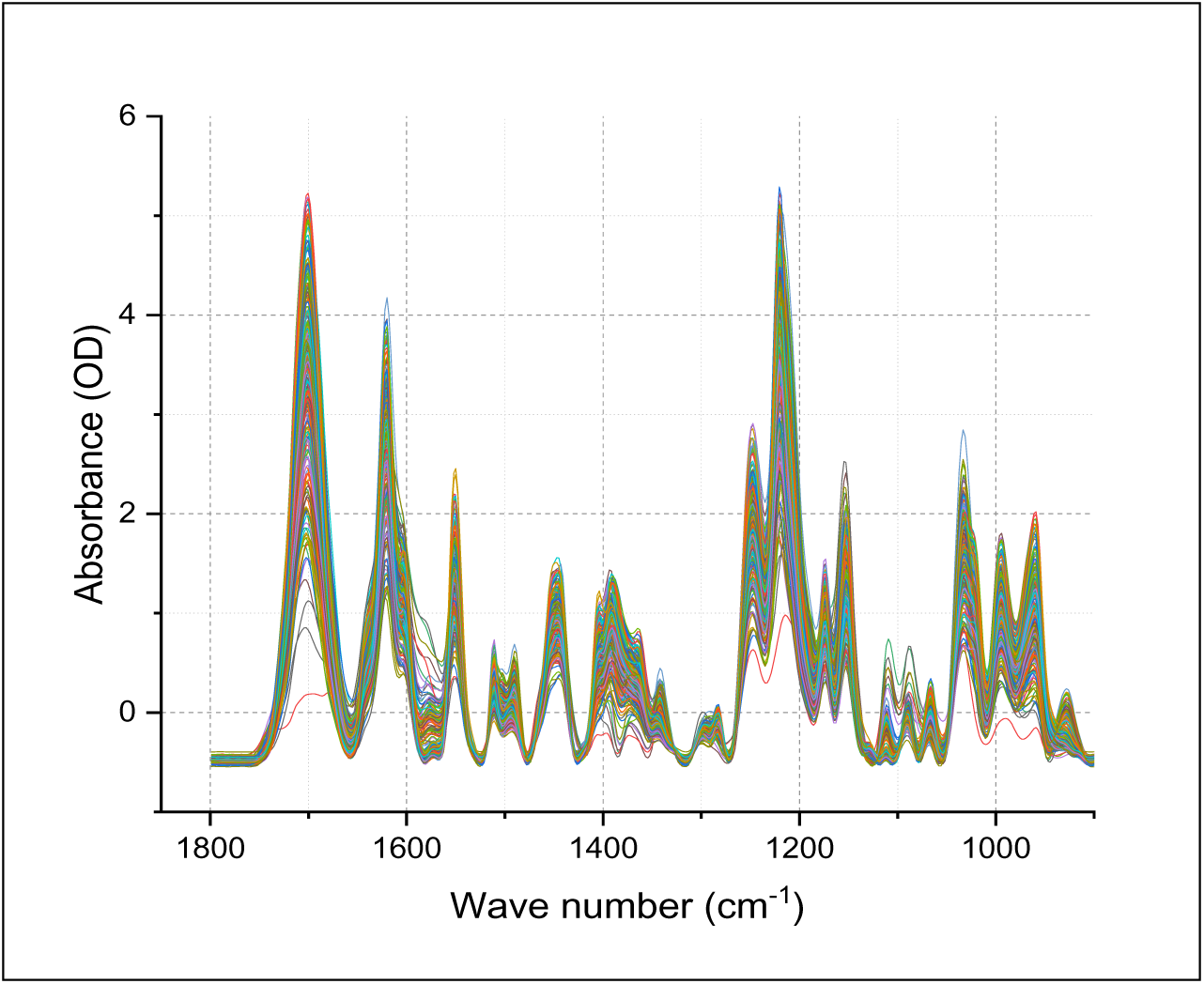
FTIR Spectral distribution of dried kava extracts from Malo and Santo Island, Vanuatu. Both noble and tudei samples were displayed.

The spectral assignments outlined in Table 1, highlight clear differences between noble and tudei/wild kava cultivars. Noble kavas are characterized by aromatic C=C bands (1600–1500 cm⁻¹) associated with yangonin and desmethoxyyangonin, as well as C–H bending (1450–1400 cm⁻¹) link to kavain. The out-of-plane C=C–H vibrations (980–950 cm⁻¹) from Methysticin serves as a distinct marker. In contrast, tudei and wild kavas show distinctive conjugated C=O stretching at ∼1650 cm⁻¹ and strong C–O stretching bands in the 1250–1000 cm⁻¹ region, corresponding to DHK and DHM. These marker peaks provide reliable spectral signatures for distinguishing between noble and tudei cultivars.

Figure 5 A and B illustrates the detection of adulteration in noble kava products through gradual substitution (w/w) with tudei and wichmannii kava cultivar. The Fourier-transform infrared (FTIR) spectrum of pure noble kava extract shows a baseline absorbance of 0 at the 1585 cm⁻¹ region, with a corresponding colorimetric optical density (OD) of 0.35. As the level of adulteration increases, the FTIR absorbance at 1580 cm⁻¹ rises progressively above the baseline. Notably, even a 1% substitution with tudei kava or wild kava results in a measurable increase in FTIR absorbance at this spectral region. Similarly, the colorimetric OD increases proportionally with higher levels of adulteration. At 30% adulteration, the OD reaches 0.83, approaching the upper limit of the nobility threshold (0.84 ± 0.05), while at 40% adulteration, the OD exceeds the threshold, reaching 0.97. The adulteration of noble kava with wichmannii cultivars showed FTIR detection of fraud at 1% substitution however, the colorimetric absorbance can only detect change in absorbance above the noble threshold with over 10% substitution. These results demonstrate that FTIR spectroscopy can detect adulteration at levels as low as 1% or less, making it a more sensitive and advanced method compared to the colorimetric assay for identifying the presence of non-noble (tudei) or wichmannii kava cultivars.

**Figure 5.**
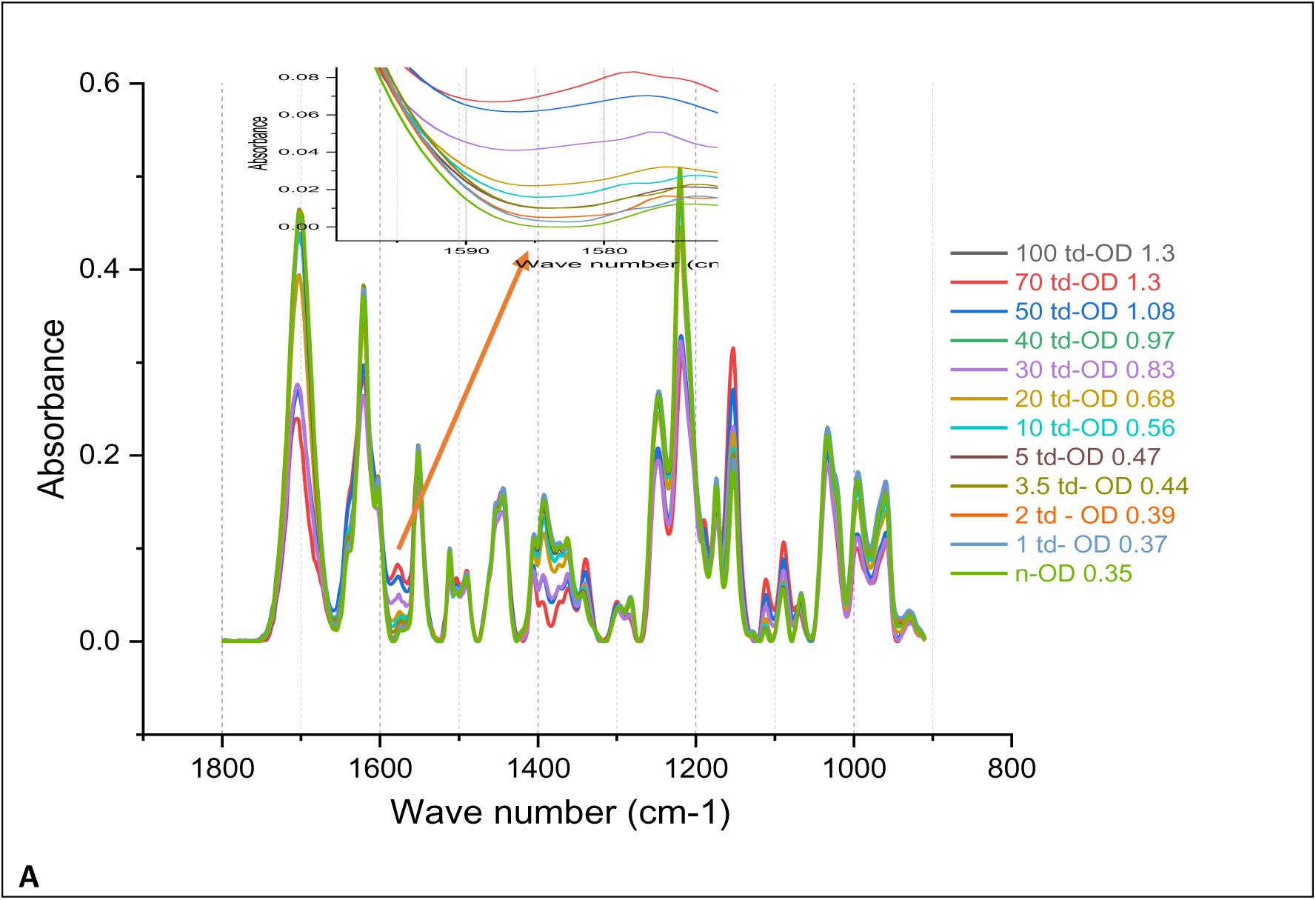
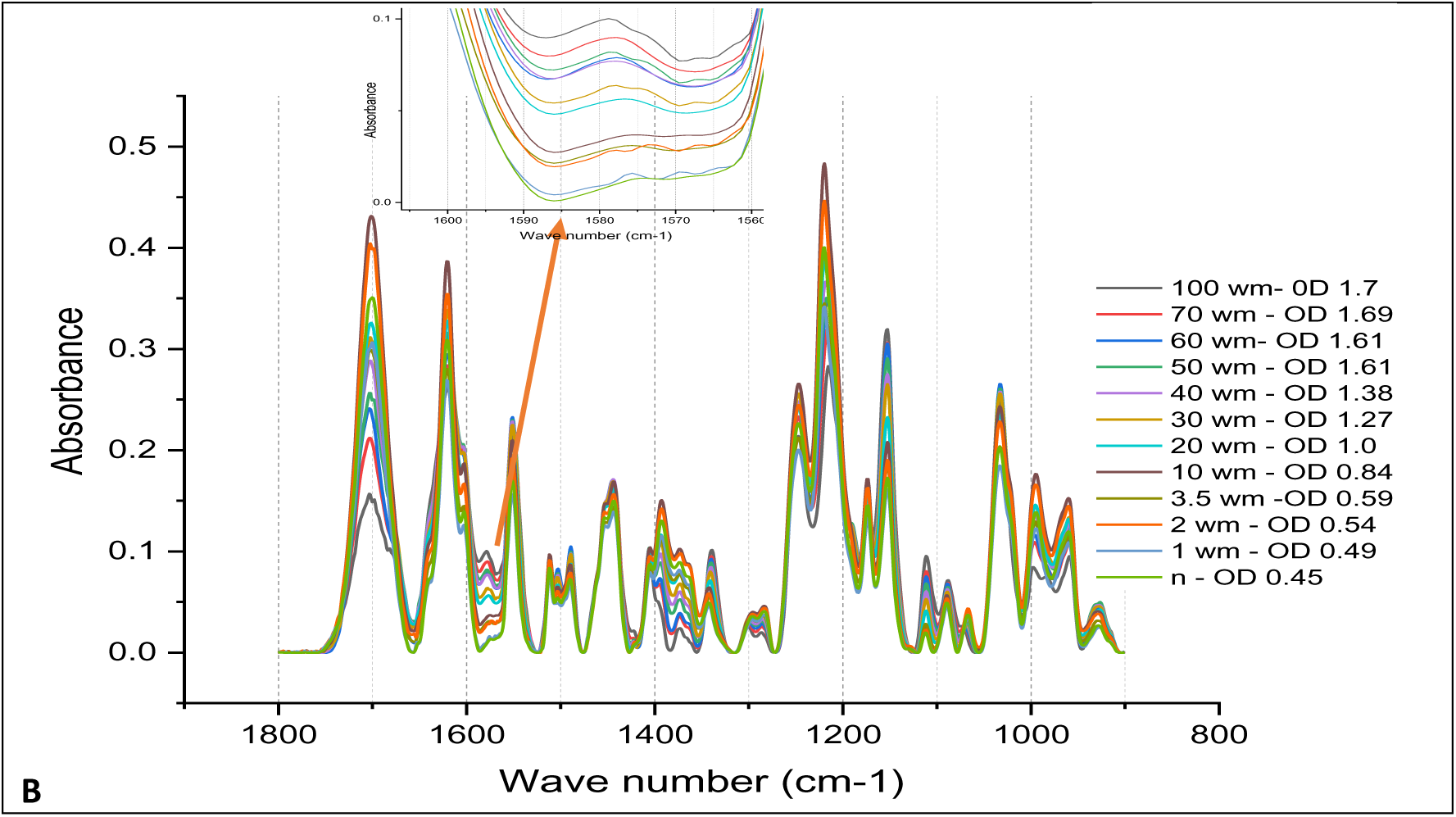
**A**, The adulteration of noble kava with tudei cultivar and, **B**., adulteration of noble kava wild kava cultivars. The numbers (0-100) represent percent (%) adulteration. The td, n and wm represent tudei, noble and wichmannii (wild kava) cultivars respectively. The OD represent colorimetric absorbance. Arrow showed 1585 region of visual differentiation.

### 3.2 ANN Classification Performance

#### 3.2.1 Geographical Origin

The artificial neural network (ANN) model demonstrated excellent predictive performance across training, validation, and test sets (Table 2). For the training set, the model achieved near-perfect fit with a generalized *R^2^* of 0.99, RMSE of 0.006, and zero misclassification. Validation results indicated strong generalization with a generalized R^2^ of 0.84, RMSE of 0.23, and a misclassification rate of 15.7%. Test set performance was also robust, with a generalized R^2^ of 0.95, root mean square error (RMSE) of 0.23, and a misclassification rate of 7.7%. Class-wise accuracy analysis revealed perfect classification (100%) for Hawaii (HI), Papua New Guinea (PG), and Vanuatu (VU) across all datasets, while Fiji (FJ) showed reduced accuracy, achieving 33.3% in validation and 0% in the test set. For Fiji, the model achieved 100 % classification accuracy under repeated-measure validation (data not presented); however, the “0 % test accuracy” shown in Table 2 does not represent a true blind-test result. No independent Fiji samples were available for external validation, and therefore the model could not be evaluated on unseen data for that region. The apparent zero accuracy reflects the absence of Fiji samples in the test subset rather than an actual misclassification failure.

**Table 2.** ANN performance on region of origin of dried kava samples.

| **Metric / Class** | **Training** |  |  |  | **Validation** |  |  |  | **Test** |  |  |  |
| --- | --- | --- | --- | --- | --- | --- | --- | --- | --- | --- | --- | --- |
| **Generalized R<sup>2</sup>** | 0.99 |  |  |  | 0.84 |  |  |  | 0.95 |  |  |  |
| **Entropy R<sup>2</sup>** | 0.99 |  |  |  | 0.72 |  |  |  | 0.85 |  |  |  |
| **RMSE** | 0.01 |  |  |  | 0.23 |  |  |  | 0.23 |  |  |  |
| **Mean Abs Dev** | 0.003 |  |  |  | 0.07 |  |  |  | 0.09 |  |  |  |
| **Misclassification Rate** | 0.00 |  |  |  | 0.06 |  |  |  | 0.07 |  |  |  |
| **Sample Size (n)** | 90 |  |  |  | 35 |  |  |  | 13 |  |  |  |
| **Class** | **Training accuracy (%)** |  |  |  | **Validation accuracy (%)** |  |  |  | **Test accuracy (%)** |  |  |  |
| **FJ** | 100 |  |  |  | 33.3 |  |  |  | 0 |  |  |  |
| **HI** | 100 |  |  |  | 100 |  |  |  | 100 |  |  |  |
| **PG** | 100 |  |  |  | 100 |  |  |  | 100 |  |  |  |
| **VU** | 100 |  |  |  | 100 |  |  |  | 100 |  |  |  |
| **Confusion Matrix Counts** | **Train predicted** |  |  |  | **Validation predicted** |  |  |  | **Test predicted** |  |  |  |
| **Actual** | FJ | HI | PG | VU | FJ | HI | PG | VU | FJ | HI | PG | VU |
| FJ | 1 | 0 | 0 | 0 | 1 | 1 | 0 | 1 | 0 | 0 | 0 | 1 |
| HI | 0 | 5 | 0 | 0 | 0 | 1 | 0 | 0 | 0 | 2 | 0 | 0 |
| PG | 0 | 0 | 9 | 0 | 0 | 0 | 3 | 0 | 0 | 0 | 2 | 0 |
| VU | 0 | 0 | 0 | 75 | 0 | 0 | 0 | 28 | 0 | 0 | 0 | 8 |

❖ FJ – Fiji ; HI – Hawaii ; PG – Papua New Guinea ; VU – Vanuatu

#### 3.2.2 Noble and tudei cultivars in regions of origin

The Artificial Neural Network (ANN) model demonstrated excellent performance in differentiating noble and tudei kava cultivars across regions (Table 3). The model achieved high goodness-of-fit, with Generalized R² values of 0.95, 0.96, and 0.97 for training, validation, and test sets, respectively, and Entropy R² values ranging from 0.90 to 0.95, indicating strong predictive capability. Error metrics were low, with RMSE decreasing from 0.14 in training to 0.07 in the test set and Mean Absolute Deviation (MAD) from 0.04 to 0.03, while misclassification rates were minimal (2% in training and 0% in validation and test). Class-wise accuracy further confirmed robustness across regions: noble kava (n) was correctly classified with 97.87–100% accuracy, and tudei kava (td) with 97.67–100% accuracy. Confusion matrices supported these results, showing very few misclassifications in the training set and perfect classification in validation and test sets. These findings indicate that the ANN model is a highly reliable tool for distinguishing kava cultivars and verifying their geographic origin, making it suitable for quality control and authentication purposes.

**Table 3.** ANN performance on noble and tudei in whole region.

| Metric/Class | Training |  | Validation |  | Test |  |
| --- | --- | --- | --- | --- | --- | --- |
| **Generalized $R^2$** | 0.95 |  | 0.96 |  | 0.97 |  |
| **Entropy $R^2$** | 0.90 |  | 0.91 |  | 0.95 |  |
| **RMSE** | 0.14 |  | 0.13 |  | 0.07 |  |
| **Mean Absolute Deviation** | 0.04 |  | 0.05 |  | 0.03 |  |
| **Misclassification Rate** | 0.02 |  | 0.00 |  | 0.00 |  |
| **-Loglikelihood** | 6.40 |  | 2.05 |  | 0.42 |  |
| **Sample Size (n)** | 90 |  | 35 |  | 13 |  |
| **Class** | **Training accuracy %** |  | **Validation Accuracy %** |  | **Test Accuracy %** |  |
| **n** | 97.87 |  | 100 |  | 100 |  |
| **td** | 97.67 |  | 100 |  | 100 |  |
| **Confusion Matrix Counts** | **Train predicted** |  | **Validation predicted** |  | **Test predicted** |  |
| **Actuals** | **n** | **t** | **n** | **t** | **n** | **T** |
| **n** | 46 | 1 | 15 | 0 | 9 | 0 |
| **td** | 1 | 42 | 0 | 20 | 0 | 4 |

❖ n – noble ; td - tudei

#### 3.2.3 Classification of quality by dried kava extract

Figure 6 shows PCA classification of noble and tudei kava using dried kava sample. The plot indicated that PCA can only distinguished between these two cultivars explaining 73.4 % of the variances in component 1 and 2, indicating the complexity of the chemical composition of the matrix.

**Figure 6.**
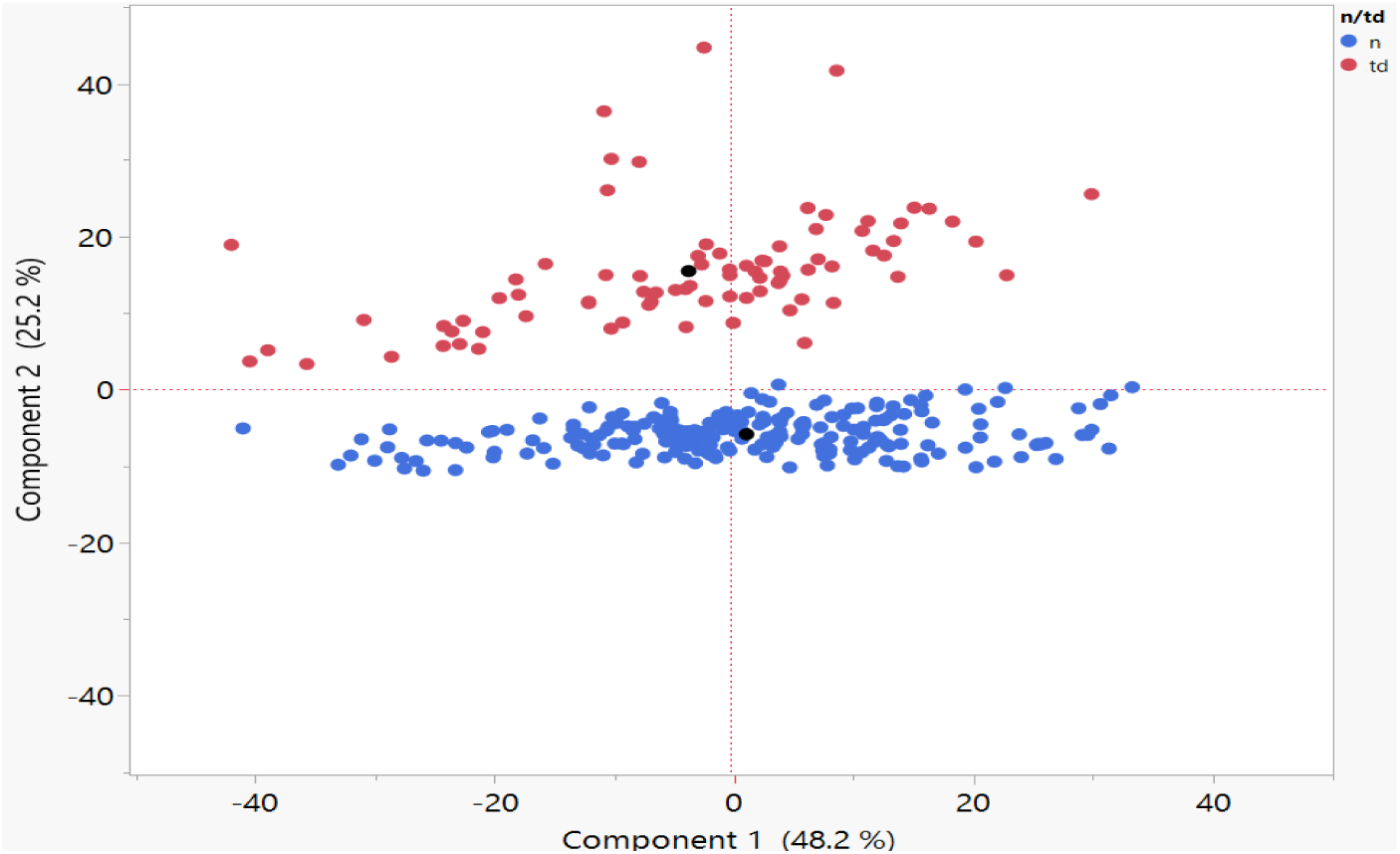
Principle component plot of noble and tudei cultivars of kava using dried kava extract. The blue points represent the noble (n) groups while the red represent tudei (td) groups.

The use of Artificial Neural Network (NTanH (3)) exhibited excellent performance in classifying kava quality classes based on dried kava extract spectra. The model demonstrated near-perfect predictive power, with Generalized R² and Entropy R² values of 1.00 for training and validation, and 0.99 for the test set (Table 4). Error measures were negligible, with RMSE and mean absolute deviation approaching zero for both training and validation, and 0.001 in the test set. Misclassification rates were 0% across all datasets, indicating that all samples were classified correctly.

**Table 4.** ANN classification of kava quality (noble/tudei) using dried kava extract.

| **Metric / Class** | **Training** |  | **Validation** |  | **Test** |  |
| --- | --- | --- | --- | --- | --- | --- |
| **Generalized R<sup>2</sup>** | 1.00 |  | 1.00 |  | 0.99 |  |
| **Entropy R<sup>2</sup>** | 1.00 |  | 1.00 |  | 0.99 |  |
| **RMSE** | $1.99 \times 10^{-7}$ | | $1.67 \times 10^{-8}$ | | 0.001 | |
| **Mean Absolute Deviation** | $2.32 \times 10^{-8}$ | | $4.15 \times 10^{-9}$ | | 0.001 | |
| **Misclassification Rate** | 0 |  | 0 |  | 0 |  |
| **-Log likelihood** | $4.17 \times 10^{-6}$ | | $3.11 \times 10^{-7}$ | | 0.01 | |
| **Sample Size (n)** | 180 |  | 75 |  | 45 |  |
| **Class** | **Training Accuracy %** |  | **Validation Accuracy %** |  | **Test Accuracy %** |  |
| **n** | 100 |  | 100 |  | 100 |  |
| **td** | 100 |  | 100 |  | 100 |  |
| **Confusion Matrix** | **Train predicted** |  | **Validation predicted** |  | **Test predicted** |  |
| **Actuals** | **n** | **td** | **n** | **td** | **n** | **td** |
| **n** | 128 | 0 | 54 | 0 | 36 | 0 |
| **td** | 0 | 52 | 0 | 21 | 0 | 9 |

❖ n- noble ; td - tudei

Class-specific accuracy confirmed the robustness of the model. Both noble (n) and tudei (td) classes were classified with 100% accuracy in training and validation, and noble also achieved 100% in the test set. Tudei maintained a perfect classification in training and validation, with all test samples correctly identified, except for the slightly smaller sample size in the test set (n = 9). The confusion matrix analysis further reinforced these findings. In the training and validation sets, there were no misclassifications between noble and tudei kava. In the test set, all noble samples were accurately classified, and all tudei samples were correctly identified.

#### 3.2.4 ANN classification of kava cultivars using dried kava extract

The neural network model (NTanH with 5-fold cross-validation) shown in table 5, demonstrated strong predictive performance with high generalized R2 values for both training (0.92) and validation (0.94) datasets, indicating an excellent model fit. Error metrics were low, with RMSE of 0.45 for training and 0.38 for validation, and misclassification rates of 21.1% and 20.0%, respectively, suggesting minimal overfitting. Class-wise analysis showed high accuracy for most classes (a: 78–80%, b: 85–87%, k: 88–100%, l: 71–80%, m: 80–88%), while class p exhibited lower accuracy (42–50%), indicating greater classification difficulty for this group. Overall, the model achieved consistent performance across folds, supporting its reliability for classification tasks.

**Table 5.** ANN classification of kava cultivars using dried kava extract.

| Metric / Class | Training |  |  |  |  |  | Validation |  |  |  |  |  |
| --- | --- | --- | --- | --- | --- | --- | --- | --- | --- | --- | --- | --- |
| **Generalized RSquare** | 0.92 |  |  |  |  |  | 0.94 |  |  |  |  |  |
| **Entropy RSquare** | 0.66 |  |  |  |  |  | 0.73 |  |  |  |  |  |
| **RMSE** | 0.42 |  |  |  |  |  | 0.38 |  |  |  |  |  |
| **Mean Abs Dev** | 0.32 |  |  |  |  |  | 0.30 |  |  |  |  |  |
| **Misclassification Rate** | 0.21 |  |  |  |  |  | 0.20 |  |  |  |  |  |
| **-log likelihood** | 121.31 |  |  |  |  |  | 23.68 |  |  |  |  |  |
| **Sample Size** | 223 |  |  |  |  |  | 55 |  |  |  |  |  |
| **Class** | **Accuracy (%)** |  |  |  |  |  | **Accuracy (%)** |  |  |  |  |  |
| **a** | 78.3 |  |  |  |  |  | 80.0 |  |  |  |  |  |
| **b** | 85.0 |  |  |  |  |  | 86.7 |  |  |  |  |  |
| **k** | 87.5 |  |  |  |  |  | 100 |  |  |  |  |  |
| **l** | 80.0 |  |  |  |  |  | 71.4 |  |  |  |  |  |
| **m** | 88.2 |  |  |  |  |  | 80.0 |  |  |  |  |  |
| **p** | 41.9 |  |  |  |  |  | 50.0 |  |  |  |  |  |
| **Confusion matrix** | **Training predicted** |  |  |  |  |  | **Validation predicted** |  |  |  |  |  |
| **Actuals** | **a** | **b** | **k** | **l** | **m** | **p** | **a** | **b** | **k** | **l** | **m** | **P** |
| **a** | 4 | 0 | 0 | 1 | 0 | 0 | 4 | 0 | 0 | 0 | 0 | 0 |
| **b** | 0 | 13 | 0 | 0 | 2 | 0 | 0 | 13 | 0 | 0 | 2 | 0 |
| **k** | 0 | 0 | 2 | 0 | 0 | 0 | 0 | 0 | 2 | 1 | 0 | 0 |
| **l** | 2 | 0 | 0 | 5 | 0 | 0 | 2 | 0 | 0 | 5 | 0 | 0 |
| **m** | 0 | 1 | 0 | 0 | 16 | 1 | 0 | 1 | 0 | 0 | 16 | 1 |
| **p** | 0 | 2 | 0 | 0 | 2 | 4 | 0 | 2 | 0 | 0 | 2 | 4 |

❖ a – ambae (bili); b – borogu ; k – klis hand ; l – palisi ; m – melomelo ; p - palarasul

#### 3.2.5 ANN classification of kava cultivars and quality using fresh kava extract

The artificial neural network (ANN) model (NTanH5) with five hidden units using k-fold cross validation as shown in table 6, demonstrated good overall performance in classifying the samples. For the training set, the model achieved a generalized R² of 0.86 and an entropy R² of 0.54, indicating a strong relationship between the predicted and observed classes. The root mean square error (RMSE) was 0.50, and the misclassification rate was 28.45%. Validation results were slightly better, with a generalized R² of 0.91, entropy R² of 0.62, RMSE of 0.46, and a misclassification rate of 20.69%.

**Table 6.**
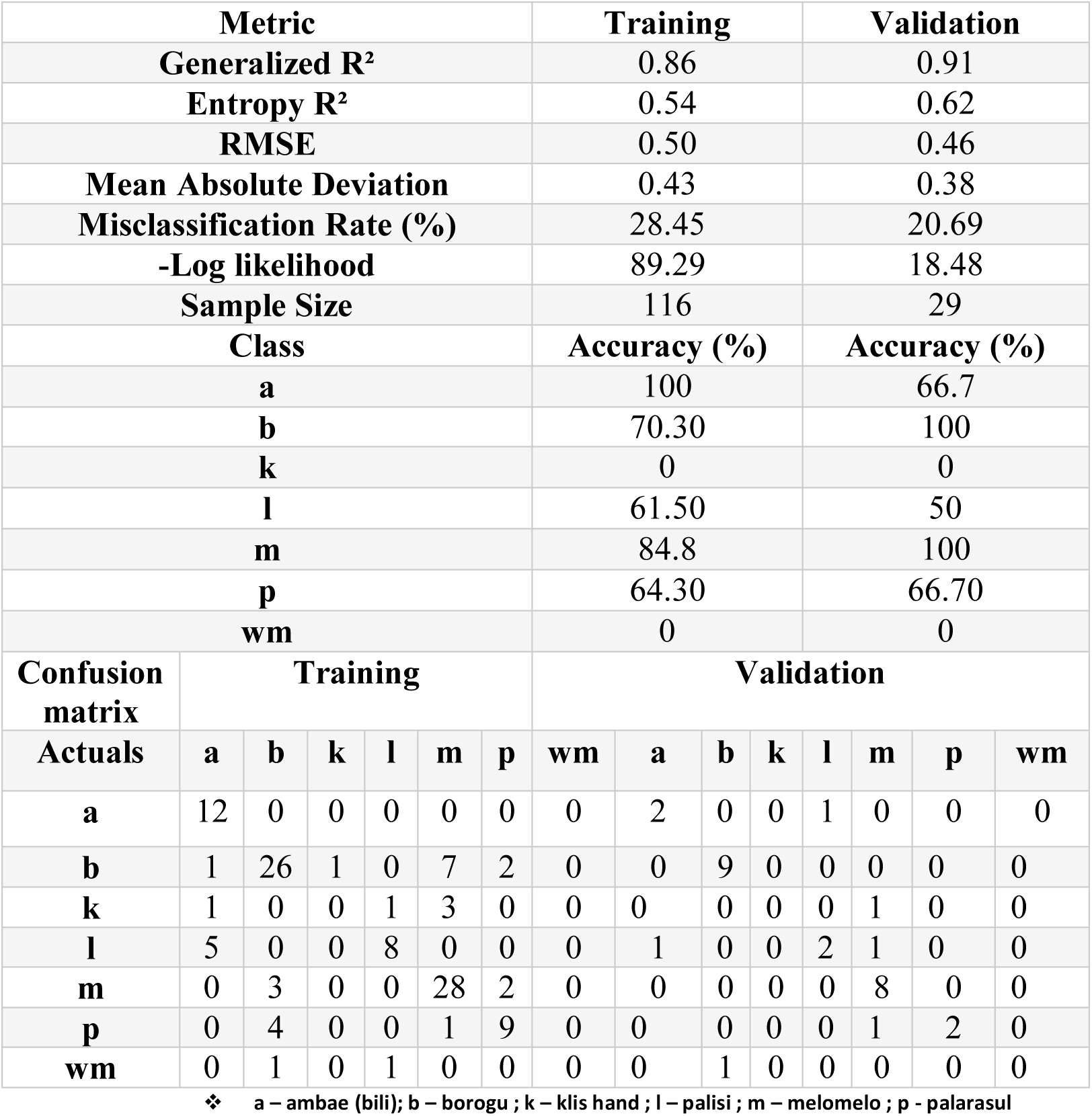
ANN performance metrics for kava cultivars using fresh kava extract.

Class-wise analysis revealed variability in predictive performance across different classes. In the training set, class a, and m were classified with high accuracy (100% and 84.8%, respectively), whereas classes k and wm were poorly predicted, with no correct classifications. Moderate performance was observed for class b (70.3%), l (61.5%), and p (64.3%). The validation set followed a similar trend, with perfect classification for class b and m, but poor performance for k and wm. Some confusion was observed between class a, and l, as well as between b and m, indicating overlaps in their feature patterns.

The Artificial Neural Network model (NTanH (3)) exhibited strong predictive performance in classifying kava quality categories based on fresh extract spectra. The model achieved high goodness-of-fit values, with Generalized R² of 0.96 (training), 0.98 (validation), and 0.96 (test), and corresponding Entropy R² values of 0.89, 0.93, and 0.92, respectively (Table 7). Error measures were low, with RMSE ranging from 0.17 (training) to 0.13 (test), and mean absolute deviation below 0.06 across all sets. Misclassification rates were minimal, at 3.45% for training, 2.78% for validation, and 4.55% for test set, confirming the strong model generalization capability. Class-level performance analysis (Table 7) indicated that noble kava (n) and suspected tudei (st) were classified with near-perfect accuracy across all datasets. Noble kava achieved 98.2% accuracy in training and 100% accuracy in both validation and test sets, while suspected tudei was correctly classified with 100% accuracy across all sets. Tudei (td) showed slightly lower accuracy, particularly in the test set (83.3%), due to occasional confusion with wichmannii (wm). The lowest performance was observed for wichmannii, which was correctly classified in only 50% of training samples, 100% in validation, but misclassified entirely in the test set. Confusion matrix analysis revealed that noble and suspected tudei categories exhibited robust separation from other classes, while wichmannii and tudei showed overlapping profiles, leading to misclassification.

**Table 7.**
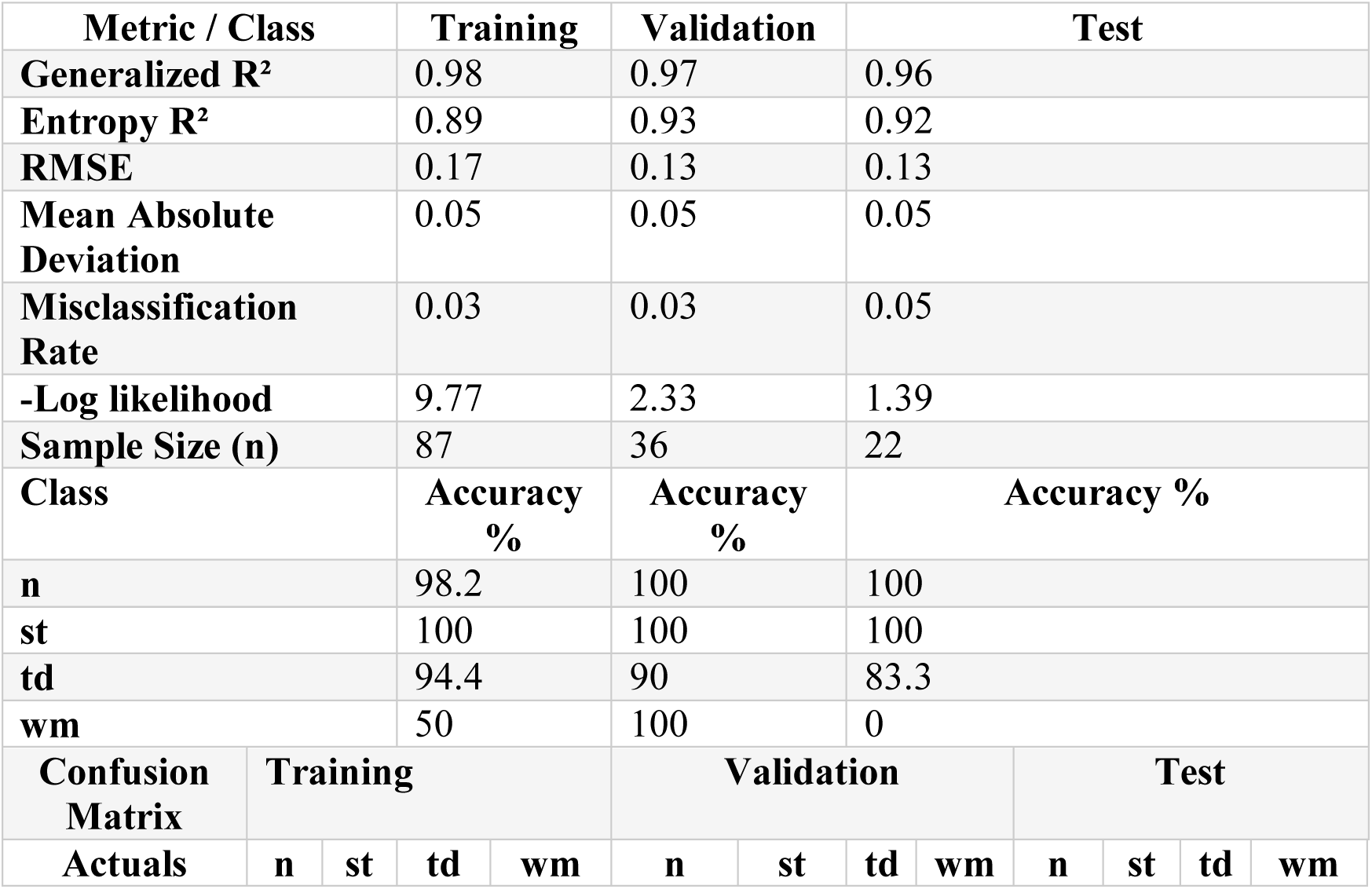
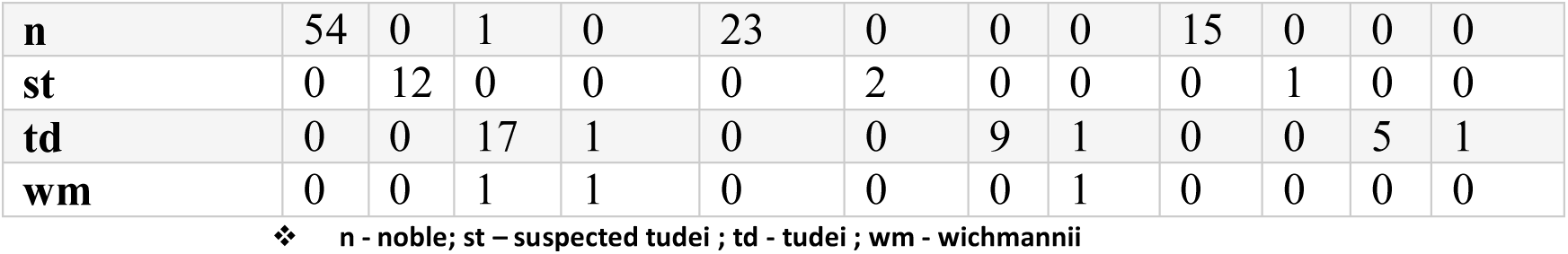
ANN performance metrics for kava quality (noble/tudei/suspected tudei/wichmannii) using fresh kava extract.

#### 3.2.6 LDA classification of kava cultivars and quality with fresh extract

The 4-class LDA model provided near-perfect discrimination (99.3% accuracy, Entropy RSquare = 0.992), with only one misclassification between noble and tudei cultivars (Table 8). In contrast, the 6-class LDA model achieved lower performance (82.8% accuracy, Entropy RSquare = 0.711), with 25 misclassifications, particularly between closely related classes (e.g., a, and l, b and m). These results suggest that classification is more robust when kava cultivars are grouped into broader chemotypic categories, while finer sub-class differentiation introduces greater overlap and reduces predictive power.

**Table 8.** LDA performance metrics for fresh kava acetone extract classification.

| **Metric** | **4-Class Model (n, st, td, wm)** | **6-Class Model (a, b, k, l, m, p, wm)** |
| --- | --- | --- |
| **Total Samples (N)** | 145 | 145 |
| **Correctly Classified** | 144 | 120 |
| **Misclassified** | 1 | 25 |
| **Overall Accuracy (%)** | **99.3** | **82.8** |
| **Percent Misclassified (%)** | 0.69 | 17.24 |
| **Entropy RSquare** | 0.992 | 0.711 |
| **-2 Log Likelihood** | 2.17 | 139.79 |
| **Best Performing Class(es)** | st, td, wm (100%) | k, wm (100%) |
| **Lowest Performing Class(es)** | n (98.9%) | a (76.5%), m (82.9%) |

❖ n - noble; st – suspected tudei ; td - tudei ; wm - wichmannii

Figures 7A and 7B present the LDA classification of noble (n) and tudei (td) kava cultivars, as well as the seven (7) cultivars. Suspected tudei (st) samples and *wichmannii* (wm) exhibited distinct chemical profiles compared with cultivated cultivars. Clear separation between noble and tudei groups was achieved based on the ATR-FTIR chemical fingerprints of fresh acetone extracts, highlighting the discriminatory power of the method. In figure 7B, some samples of Palarasul (p) and Borogu (b) and Klis hand (k) were in the suspected tudei group due to their unique chemical profile. The wichmannii (wm) and tudei (td) cultivars like Ambae Bili (a), palisi (l) and Klis hand (k) were clearly isolated to their respective classes. The famous noble kava groups with red points are the Melomelo (m), Borogu (b) and Palarasul (p).

**Figure 7.**
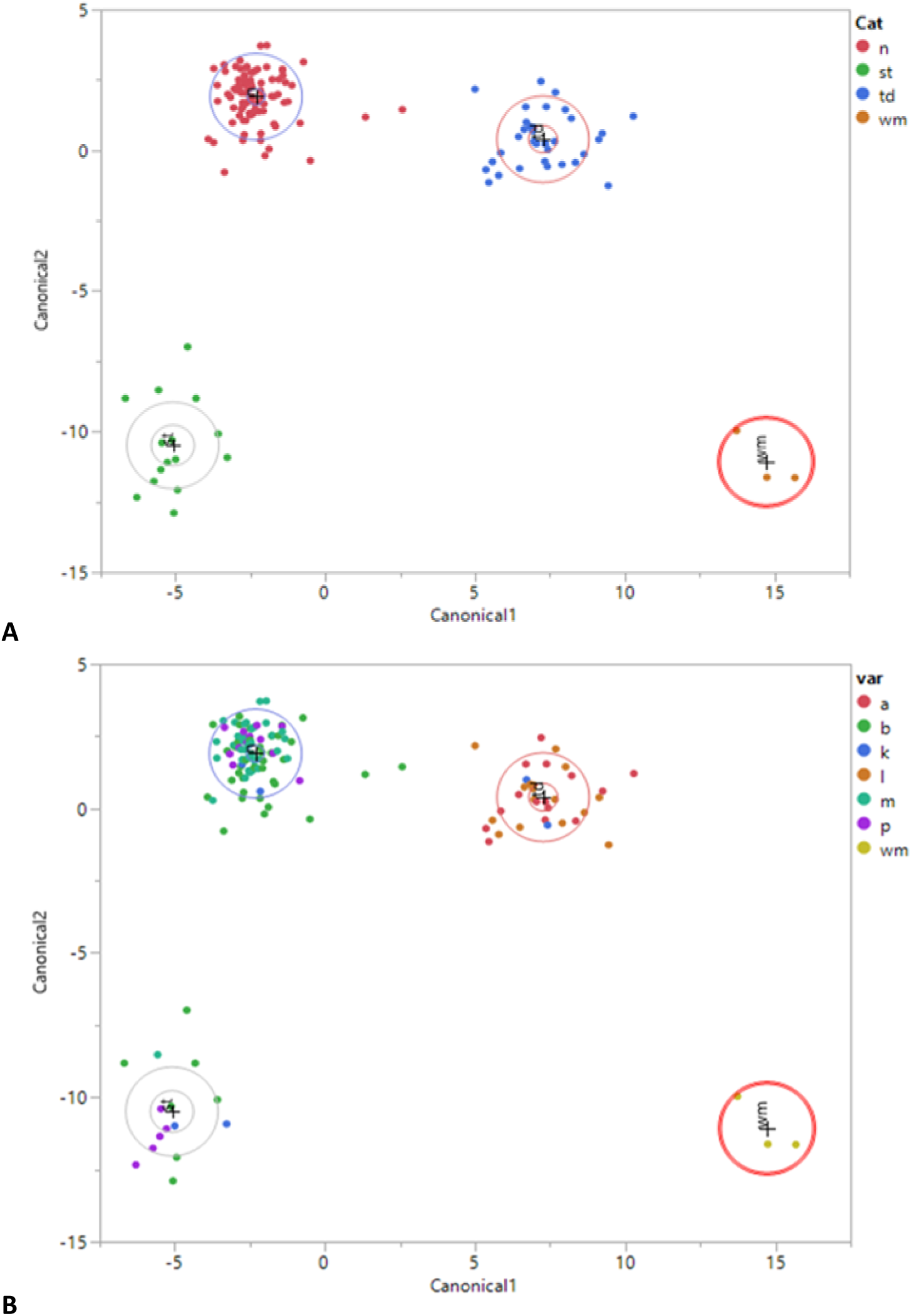
LDA classification of Kava by cultivars and quality (noble/tudei). A, represent the 4 kava category: n-noble, st-suspected tudei, td-tudei and wm-whichmanii. B, present 7 cultivars with local names: a – Bili (ambae), b – Borogu (Red hand), k – Klis hand, l – palisi, m – melomelo (Green hand), p – palarasul (Yellow leaf), and wm (wichmannii).

#### 3.2.7 ANN classification of kava by island of origin

The ANN model (NTanH5 with 5 hidden layers (k-fold cross validation)) achieved perfect classification accuracy (100%) for both training and validation sets, with zero misclassifications across all classes (m and s) (Table 9). The high generalized *R^2^* values (0.92 for training and 0.91 for validation), combined with very low RMSE and Mean Absolute Deviation, indicate excellent model fit and no signs of overfitting. These results suggest that the spectral characteristics of kava from islands m (Malo) and s (Santo) are highly distinct, making classification straightforward for the ANN model.

**Table 9.** ANN performance metrics on kava samples classification with island of origin.

| Metric / Class | Training |  | Validation |  |
| --- | --- | --- | --- | --- |
| **Generalized $R^2$** | 0.92 |  | 0.91 |  |
| **Entropy $R^2$** | 0.89 |  | 0.88 |  |
| **RMSE** | 0.08 |  | 0.06 |  |
| **Mean Abs Dev** | 0.028 |  | 0.027 |  |
| **Misclassification Rate** | 0.00 |  | 0.00 |  |
| **Sample Size** | 116 |  | 29 |  |
| **Class (Island)** | **Accuracy (%)** |  | **Accuracy (%)** |  |
| **m** | 100 |  | 100 |  |
| **s** | 100 |  | 100 |  |
| **Confusion Matrix** | **Training (m / s)** |  | **Validation (m / s)** |  |
| **Actuals** | m | s | m | s |
| **m** | 10 | 0 | 2 | 0 |
| **s** | 0 | 106 | 0 | 27 |

❖ m - malo; s - santo

### 4.0 Discussion

This study demonstrates the effectiveness of Artificial Neural Networks (ANN) combined with spectral analysis for authenticating the geographical origin, cultivars, and quality of kava (*Piper methysticum*). Overall, the ANN models achieved high predictive accuracy, low error rates, and strong generalization across multiple classification tasks, underscoring their potential as robust tools for quality assurance and traceability in the kava industry.

### 4.1 Geographical Origin Authentication

The ANN model (TanH3) successfully classified kava samples according to their countries of origin, achieving a generalized R² of 0.99 (training), 0.84 (validation), and 0.95 (test). Class-specific accuracy was perfect (100%) for Hawaii (HI), Papua New Guinea (PG), and Vanuatu (VU), while Fiji (FJ) showed lower accuracy (33.3% validation, 0% test). This reduced performance suggests Fijian kava shares overlapping chemical signatures with other origins, likely due to shared cultivars, hybridization, or similar environmental conditions (2, 23). However, the ANN model with five TanH neurons and 5-fold cross-validation achieved a classification accuracy of 100% for Pacific region kava samples although the detailed performance metrics are not presented here surpassing the performance of PLSDA predictive modelling accuracy of 96.7% (5). This result highlights the model’s strong predictive capability and aligns with previous studies demonstrating the utility of ANNs for food and plant authentication (14, 20, 21). The high accuracy observed suggests that spectral features captured from kava root material are highly distinctive for regional origin differentiation Significantly, the model achieved perfect classification (100% accuracy) for Malo and Santo Island kava, demonstrating its capability to differentiate micro-regional origins within Vanuatu. Such high-resolution differentiation is critical for export certification, geographical indication labeling, and market value optimization. Similar approaches are applied in other high-value commodities, such as PDO wine and specialty coffee, where micro-terroir influences market differentiation (24, 25).

### 4.2 Differentiation of Noble vs. Tudei

ANN achieved near-perfect classification of noble and tudei cultivars across pacific island regions, with negligible misclassification using dried kava samples. These results confirm that noble and tudei kava cultivars possess distinct spectral fingerprints, primarily due to differences in kavalactone composition and flavokavain content (3, 26). Accurate identification of noble kava supports Vanuatu’s export policy, ensuring consumer safety and regulatory compliance (27).

### 4.3 Classification Based on dried and Fresh Extracts

Models based on dried kava extracts demonstrated excellent performance, achieving generalized R² of up to 1.00 with zero misclassifications, reflecting the stability of spectral features in dried material. Fresh extracts were more variable, likely due to moisture interference and complex matrices (2), making dried extract-based ANN preferable for reliable commercial authentication.

ATR-FTIR analysis of fresh acetone extracts revealed clear differentiation among kava cultivars. The number of peaks and peak height could be used to calculate and determine threshold between kava cultivars of noble and tudei kava. Noble kava lacked a peak at 1100 cm⁻¹, tudei cultivars exhibited prominent DHK and DHM peaks, and suspected mixed samples showed broad, high-intensity bands at 1000–1100 cm⁻¹, indicative of elevated DHK content. These spectral differences correspond to variations in kavalactone composition, with 1050–1100 cm⁻¹ bands linked to C–O stretching and 1550–1700 cm⁻¹ to C=C vibrations (28, 29). Wichmannii displayed a distinct fingerprint between 1200 and 1100 cm-1 and 1650 cm-1 compared to other cultivars, supporting the potential of ATR-FTIR as a rapid, non-destructive quality control tool without requiring advanced machine learning or predictive modeling expertise(29, 30). Dried kava acetone extracts showed clear chemical differentiation between cultivars, as reflected by the variation in the number of peaks within the 1125–900 cm⁻¹ region: five peaks for noble, six for tudei, and four for Wichmannii. These distinct spectral features indicate strong potential for accurate classification of kava cultivars without relying on predictive modeling or advanced machine learning. Furthermore, both fresh and dried kava spectra exhibited consistent cultivar differences, underscoring their potential application in quality control systems. Such rapid FTIR-based screening could be employed prior to processing and export to minimize production losses and reduce risks associated with non-compliant or misclassified kava entering international markets. Using these features, the ANN achieved 99% classification accuracy with a single hidden layer of five TanH neurons and 5-fold cross-validation. LDA also effectively separated tudei and wichmannii from noble cultivars. Some samples (e.g., borogu, palarasul, klish hand) formed distinct clusters, potentially due to unique chemical profiles or nomenclature inconsistencies, yet ANN performance remained robust. Overall, these results demonstrate the combined utility of ATR-FTIR and ANN modeling for accurate kava cultivar authentication, consistent with prior work in food and plant authentication (14, 20, 21).

### 4.4 Comparison with Food Authentication Systems

The authentication of kava origin and cultivars parallels traceability systems in other high-value commodities, including wine, coffee, and herbal medicines, where spectroscopy coupled with machine learning is the standard practice (31). ANN models are particularly effective at capturing non-linear relationships in complex spectral datasets, outperforming PCA or LDA in resolving overlapping chemical signatures (32, 33).

### 4.5 Regulatory and Export Certification Implications

The ability to authenticate kava from specific islands such as Malo and Santo enable premium labeling, traceability, and fraud prevention supporting Codex Alimentarius and ISO 22000 compliance. ANN–FTIR authentication provides a rapid, non-destructive, and reliable tool for certifying both geographical origin and cultivars, which is critical for the international kava market. The quality check of fresh kava acetone extract also helps prevent business losses for exporters as test can be performed prior to processing thereby minimizing production losses due to poor quality kava. FTIR spectroscopy also provides a much more sensitive means of detecting adulteration in noble kava compared to the traditional colorimetric assay (34). Even small amounts of substitution with tudei or wild kava (as low as 1%) produced visible and measurable spectral changes at 1585 cm⁻¹, whereas the colorimetric method only approached its detection threshold (absorbance ∼ 0.84 ± 0.05) at higher levels of adulteration (>30% for tudei and >10% for wichmannii). While the colorimetric test is still useful for quick and low-cost checks, FTIR offers a clearer and more reliable way to confirm authenticity, making it a valuable tool for protecting the integrity and market value of noble kava

### 4.6 Limitations and Future Work

Despite achieving high accuracy, classification of certain cultivars and Fijian kava remain challenging due to chemo-type overlap and limited sample size. Future work should focus on expanding spectral libraries, incorporating deep learning approaches, and combine FTIR with complementary techniques (e.g., NIR, LC-MS, HPLC) to further enhance reliability robustness of kava authentication for commercial and regulatory applications. Use of FTIR database of kava extract and calculations of spectral peak heights could enhance direct differentiation of kava quality (35).

## Conclusion

This study demonstrates the strong potential of combining ATR-FTIR spectroscopy with artificial neural network (ANN) modeling for authenticating kava origin and cultivar. The ANN achieved high classification accuracy in distinguishing noble and tudei kava cultivars, successfully differentiate kava from Pacific Island regions, and attained 100% accuracy for differentiating kava samples from the island of Malo and Santo Island in Vanuatu, highlighting its suitability for regional and micro-regional authentication. Quality checks on fresh kava acetone extracts allow early detection of tudei and wichmannii cultivars, preventing losses during processing, and visual inspection of spectral differences provides a rapid, low-cost method for fraud detection that does not require machine learning expertise. These findings support Codex Alimentarius and ISO 22000 compliance and offer a rapid, non-destructive, and cost-effective approach for food traceability, fraud prevention, and quality control in the global kava trade.

## Acknowledgements

The authors would like to acknowledge the farmers who provided kava samples and the VBS Provincial team for their assistance with sampling.

